# Mesoscale Imaging of Cortical Sensorimotor Integration in Huntington’s Disease Mice During Reward-Guided Behaviour

**DOI:** 10.1101/2025.04.23.648272

**Authors:** Kai Trappenberg, Daniel Ramandi, James Mackay, Timothy Murphy, Lynn Raymond

## Abstract

Huntington’s disease (HD) is a neurodegenerative disorder that affects numerous brain functions, yet how altered sensory processing contributes to behavioral and learning deficits remains poorly understood. Previous wide-field mesoscale imaging and electrophysiological recording from anesthetized HD mice revealed that sensory stimulation induced exaggerated, prolonged cortical activity across more brain regions compared to wildtype (WT) littermates. This suggests differences in sensory processing; as such, this study aimed to investigate the cortical activity in a cue-based sensory-guided learning task in a custom-built Raspberry Pi-controlled two-alternative forced choice (2AFC) rig. The rig was designed to enable head-fixed zQ175 knock-in HD mice and WT controls crossed with Thy1-GCaMP6s mice, to perform a cue-based visual discrimination task while mesoscale calcium imaging recorded activity across layer 2/3 of the cortex. Mice that successfully licked the reward spout displayed decreased global cortical activity before the reward presentation, with WT mice showing more spatially localized suppression than HD mice. HD mice exhibited exaggerated cortical responses to visual stimuli and prolonged cortical motor-related activity during licking. While this is an exploratory pilot study with limited sample size, preventing definitive genotype-based comparisons, the custom behavioral system lays the groundwork for future studies into how sensory processing deficits contribute to cognitive impairments in HD. This work provides an important step toward understanding the interplay between cortical circuit dysfunction and behavioral outcomes in HD, offering a novel platform to investigate early sensorimotor integration learning impairments.

## Introduction

Huntington’s disease is an inherited progressive neurodegenerative disorder caused by an expanded cytosine-adenine-guanine (CAG) repeat in the huntingtin (*HTT*) gene (1). Clinically, HD is characterized by a triad of motor, cognitive, and psychiatric symptoms (2). Emerging evidence suggests that the cognitive difficulties seen in HD may, in part, arise from disrupted sensory feedback and the brain’s reduced ability to integrate this information into motor control and cognitive planning. A growing body of research has shown sensory processing deficits in patients with HD, including auditory, tactile, and visuospatial perception, as well as cortical thinning of sensory regions (3, 4, 5). These sensory processing deficits may impair the brain’s ability to interpret feedback, hindering motor adjustments, learning, and decision-making (6, 7, 8, 9). This disrupted feedback loop could worsen motor and cognitive symptoms, making skill acquisition increasingly difficult. Given the lack of a cure, understanding these mechanisms is crucial for developing effective interventions. Brain imaging studies have revealed alterations in sensory and association cortices in HD mice, suggesting circuit-level dysfunction that may underlie these behavioral impairments (10). This aligns with other studies in HD mice, where changes in excitatory-inhibitory balance and reduced neuron-neuron correlations suggest aberrant cortical coordination and network dynamics (11, 12, 13). Given that sensorimotor integration relies on precise cortical coordination, these disruptions may impair the brain’s ability to process sensory input and translate it into appropriate motor responses. Studies in mice have shown that sensory inputs activate specific cortical areas before propagating to cortical motor and association regions, reflecting integrated processing essential for adaptive behavior (14, 15). In HD, these circuit-level disruptions may contribute to deficits in sensorimotor integration, exacerbating motor and cognitive impairments.

Recent studies in HD mouse models further support these findings, showing a more robustly exaggerated and prolonged cortical response to sensory stimuli (16). Specifically, sensory stimulation in HD models resulted in hyperexcitability and broader cortical activation, including sensory, motor, and association areas, suggesting abnormal coordination of cortical networks. These changes may underlie deficits in sensory processing and contribute to impairments in sensory learning, as observed in HD mice during go/no-go discrimination tasks (17, 18).

These findings highlight the value of mesoscale imaging in studying circuit-level dysfunction and emphasize association areas and cortico-cortical pathways as key sites of disease-related changes. Since these pathways are readily activated by sensory stimuli, they provide valuable targets for investigating circuit changes in neurodegenerative disorders like HD and offer insights into how sensory-driven cortical and motor responses are disrupted in disease.

We employed simultaneous *in vivo* calcium imaging of the entire layer 2*/*3 of the cortex during a sensory-cue based discrimination associative learning task. In our pilot project, which focused on the training stages towards a two-alternative forced choice task, we observed intriguing differences in cortical circuits between HD and WT mice.

## Methods

### Animal model

zQ175 knock-in mice on a C57BL/6 background expressing GCaMP6s under a Thy1 promoter (thy1-GCaMP6s zQ175dn) were used for this study. The transgenic thy1-GCaMP6s zQ175dn mouse lines were bred in house with only GCaMP6s positive mice expressing either Q175 positive (HD) or Q175 negative (WT) used from this colony. Animal tissue was collected through ear clipping at weaning. DNA extraction and PCR analysis were used to determine the genotype of each mouse. Mice were group-housed from weaning to experimental endpoint. The mice were housed in Animal Care Systems OptiMICE cages with wood fiber bedding (NEPCO Beta Chip), a paper hut, and nesting material from birth until 7 days after cranial window surgery. Mice had free access to food and water and were kept under a 12 hour light/dark cycle (lights on at 07:00, off at 19:00). 7 days after cranial window surgery the mice were transferred into a 12 hour reverse light cycle room (dark/light cycle) with lights on at 19:00 and off at 07:00. The cages were switched to conventional “shoe-box” cages, also with wood fiber bedding, their original paper hut and nesting materials. Mice were held in the reverse light cycle room for 10 days before they were handled and habituated to the experimental rig. All experiments were carried out in accordance with the Canadian Council for Animal Care guidelines and approved by the University of British Columbia animal care committee (protocols A23-0083 and A21-0276).

### Cranial window surgery

Intact-skull cranial window surgery was conducted on the experimental mice. To do this, mice were placed under anesthesia, then an incision over the cortex was made and the skin retracted. Then, a 1 cm x 1 cm glass coverslip was applied using Metabond clear dental cement (Parkell; Product: CB Metabond). Following this, a custom titanium headbar was cemented on. The animal’s temperature was kept at 37°C via a feedback-controlled heating pad (WPI) for the duration of the surgery. Refer to the supplementary note 1 for full surgical protocols.

### Behavioural paradigm

Mice were behaviourally tested in a custom built rig using parts from Thorlabs™, Adafruit™, Luxeon™, as well as custom built and 3D printed parts (Figure 1). These were assembled and changed as the behavioural paradigm was refined for our experimental needs. Headfixed mice were water restricted, in accordance with animal care protocols, to motivate them to perform the given task. Up to 1ml of water was presented as a reward to the mice. In the event mice did not perform well enough in the task to receive the full 1ml of water, the remaining water was given to them at the end of the day, at least 2 hours after the task was last performed. In the first stage, mice were head-restricted and presented with a start cue; 4.2 seconds after this, they were presented with two spouts (left and right). If the mice licked either spout, the opposing spout would move back away from the mouse, and they received water for the remaining 2 seconds. Then, 8 seconds later, the paradigm repeated for a total of 100 trials. Mice would move onto the next stage by successfully licking 50 percent of the trials for 2 consecutive days.

**Fig. 1.**
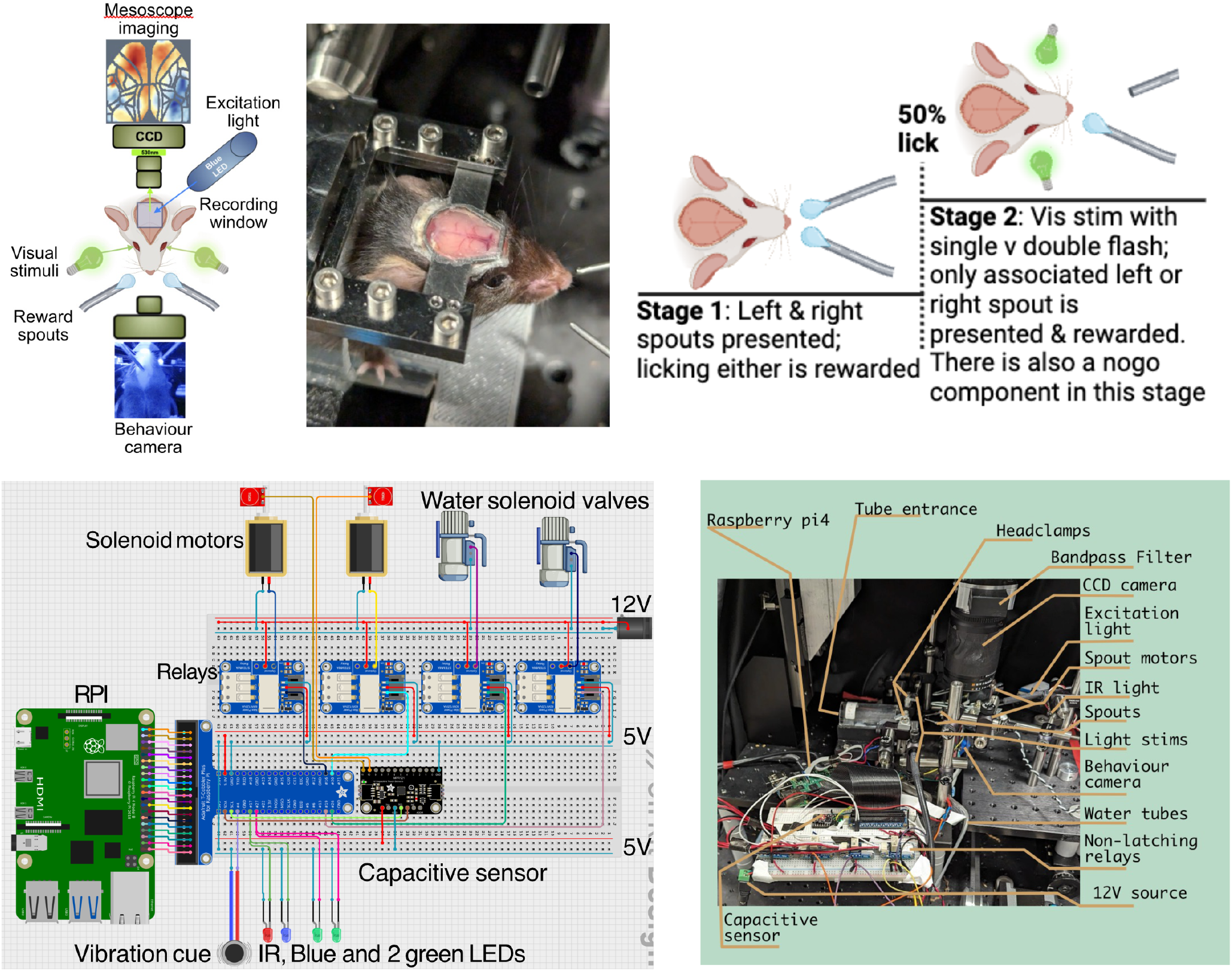
Experimental rig design. (Top Left) Mice have a cranial window with CCD recording of the cortex and behavioural camera. Complete with a blue excitation light for GCaMP, green visual stimuli and reward spouts for water. (Top Middle) Mouse cranial window and headfix apparatus. (Top Right) Behavioural training stage paradigm. Stage 1 where only spouts are presented with no visual association. Upon meeting the threshold mice move into stage 2, with a visual stimulus modality paired with a left or right water reward. (Bottom Left) Circuit diagram from the custom experimental rig. (Bottom right) Labeled experimental rig.

The second stage of learning began to include an associative component, in which a green LED light (single 120ms flash or two 60ms flashes) was paired with the presentation of the water spouts, as follows: 3 seconds after the start cue, the light flash(es) was presented, and 1.2 seconds following the light, one of the two spouts were presented. Which spout (left or right) was presented was determined by the corresponding single flash for the left spout, or two flashes separated by 1 second (double flash) paired with the right spout. Since there was only one spout presented, if the mouse licked the spout they would be rewarded. Hence this stage did not include any decision making between which spout to lick for water, and instead was intended to reinforce the pairing between the single flash with left spout versus double flash with right spout. The water would be given to the mouse for the remaining 2 seconds that the spout was presented for. This condition encompassed 100 repeated trials.

### Rig parts and assembly

The Raspberry Pi (RPi)-controlled Two-Alternative Forced Choice Task Rig was custom designed and built for this project. The following parts were used in the development and assembly of the rig. A Raspberry Pi 4 model B Quad Core 64 bit with 4GB ram (upgraded later to 8GB ram) ran the experiment using a custom python script. Additionally, a transparent plastic tube was acquired to hold the mice with dimensions 31mm inner diameter (ID), 35mm outer diameter (OD).

Custom designed parts were 3D printed on an Ender 3. This includes: the base holding the tube (which in turn holds the mouse’s body in place), a infrared camera holder, servo motor holders, a mouse paw rest (with an insert for the vibrational buzzer), and a breadboard platform. 12V dual power LED drivers were custom built by EnixLABs (Figure S1). They were used to power the green LED lights for the stimulus, the infrared light, and the blue excitation light for cortical recordings.

The headbars were custom ordered from PCBWay and cut out of titanium. These also included a custom headbar clamp. All Raspberry Pi parts were acquired from Adafruit™. The Raspberry Pi was connected to a Full Sized Premium Bread-board (830 Tie Points) via a GPIO Ribbon Cable and Cobbler Breakout Cable (see Figure 1 for image of breadboard layout and components). The breadboard contained a 4 STEMMA non-latching (defaulting to an open or closed state until actively switched, allowing 5V control signals to toggle the 12V system) mini relays, each with 2mm 3-Pin male header cables which connected to the GPIO outputs. Two of the relays were wired to two large push-pull solenoids, respectively. The other two relays were wired to a water-valve solenoid, which was fed via a 1mm tube with water in a bottle with a hole for the water tube. An MPR 121 12-key Capacitive Touch Sensor Breakout was connected to SDA and SCL on the RPi. A standard Infrared (IR) Raspberry Pi Camera Module 2 with an RPi 25mm, 5 megapixel telephoto lens and 850nm LUXEON™ IR LED (on a Saber Z1, 10mm Square Base with 1050 mW @ 1000mA). The rig connected to two Adafruit™ Micro servo (TowerPro SG92R) via general purpose input/output (GPIO). The tone start cue was generated The tone was generated by an Adafruit™ STEMMA Piezo Driver Amp at 10 KHz frequency connected to a GPIO on the RPi.

The light for visual stimulation were two Green (530nm), LUXEON™ Rebel Color LEDs (on a SABER^2^ Star Base with 102 lm @ 350mA) each attached by the same means as the excitation light (custom drilled Ø1” optical posts with theropaste and encapsulated by a ThorlabsTM C-Mount Extension Tube with two convex lenses, Thorlabs™ a Ø3 mm (LLG to SM1) Adapter to connect to a Thorlabs™Liquid Light Guide (Ø3 mm Core, 340 - 800 nm)). The cortical excitation light was a Blue (470nm) LUXEON™ Rebel LED, mounted on a 20mm Star CoolBase (70 lm @ 700mA). These were attached to custom drilled 1” diameter (Ø) optical posts with theropaste and encapsulated by a Thorlabs™ C-Mount Extension Tube with two convex lenses, and a Thorlabs™ Adapter (Ø3 mm LLG to SM1) which connects to a ThorlabsTM Liquid Light Guide (Ø3 mm Core, 340 - 800 nm).

Refer to Figure Figure 1 for an image of the rig assembly. Most parts were attached to the rig via Thorlabs components, this included Ø1/2” Optical Post in varying lengths from L = 1” to L = 6”). These posts were screwed into the baseplate or held in place with a Ø1/2” Pedestal Post Holder (Spring-Loaded Hex-Locking Thumbscrew, L=1.19”) and a Clamping Fork (1.75” Counterbored Slot). The motors and lights were attached at 90 degree angles with a Right-Angle Clamp for Ø1/2” Posts and Right-Angle Post Clamp for Ø1/2” Posts. Additionally the headbars were custom designed to screw into the Ø1/2” Optical Post.

### Imaging set up and script processing

A Monochromatic Pantera CCD Camera (Dalsa) with green emission filter (525/36nm) and front-to-front Nikon lenses (f=1.4, 35mm; f=2) was used to acquire cortical calcium data.

Each experiment (stages 1 and 2) was run with a custom python script. Refer to Github (https://github.com/ktrap06/RPI_rig) for further details on the code. These scripts generated the behavioural dataset of success rates as a txt file. The txt files were imported into a custom python script for further analysis and plotting.

The datasets generated in this project are videos of cortical surface calcium dynamics in layer 2/3 of the cortex (stored as tiff stacks in approximately 70 000 depth x 128 height x 128 width). A python pipeline was coded to process the cortical data (refer to open source Github link - github.com/ktrap06/corticalanalysis).

The preprocessing step began with removing any dark frames at the beginning and end of the experiment, followed by the identification of the green visual stimuli used to elicit cortical network activity. Since this light creates an artifact in the cortical signal, it is removed in order to calculate a change in fluorescence over the baseline fluorescence (.6.F/F). The removed frames are then interpolated using the signal before and after the artifact. Then the .6.F/F is calculated to normalize the signal to a 10 second moving average thereby filtering out any hemodynamic or broader movement artifacts. The tiff stack is then spatially smoothed with a 3 pixel gaussian blur, and temporally smoothed with a chebyshev type 1 temporal filtering (high and low bandpass filter). The database was then analyzed with heatmaps and brain panel scripts for further analysis.

Each session was imported into a master script and subdivided based on HD or WT, single or double flash, successful lick or no-lick, and by day. All data was z scored to normalize between mice. A mask was applied to each brain panel using regions from the standardised Allen Brain Atlas.

## Results

### Behavioural results

In stage 1 (Figure 2), both HD and WT groups exhibited steady improvements in success rates, with HD mice requiring more days to learn to lick, compared to WT mice. All mice in this set of experiments were female. In spite of successfully licking the spouts, in stage 2 (Figure 2), neither genotype group surpassed the success thresholds for “success lick” and “success no-go” exceeding 75 percent for two consecutive days. Both the HD and WT groups exceeded the 75 percent threshold for successful licks; however, the no-go success rates on those days were notably low. This suggests that the mice were licking on every trial without paying attention to the stimulus. These results highlight the difficulty in suppressing the incorrect licking responses, a behavior likely overtrained during stage 1. Nevertheless, mice learned to associate the flash of light with licking the spouts for water rapidly, but were not able to make the association with left or right spouts and their respective single or double flash light condition.

**Fig. 2.**
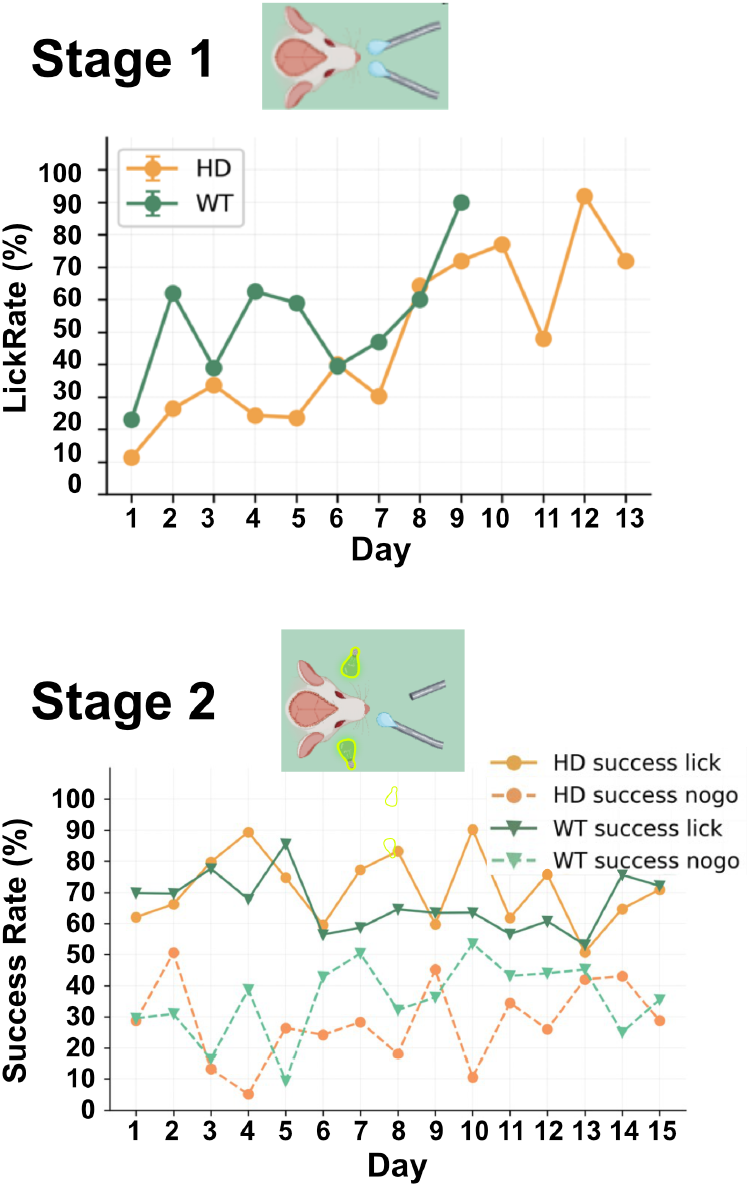
Lick rates for mic in stage 1 and 2 respectively (n=5). (Top) Mice progressed through stage 1 before reaching the licking threshold. (Bottom) Once in stage 2, mice plateaued in their performance on both left/right stim and no-go trials

### Primary visual cortex activation requires visual stimuli of duration greater than 30-ms

A set of experiments were carried out with varying visual stimulus durations. The flash stimulus parameters consisted of a 60-ms single flash and a 30-ms double flash (two flashes separated by 0.5 seconds). However, analysis of cortical activity revealed a stark difference in the neural response between these two conditions. The 30-ms double flash condition (Figure 3) produced minimal activation in the primary visual cortex (V1) across the entire 17-day experimental period in both HD and WT mice. This lack of a strong response suggested that the shorter flash duration might not be sufficiently salient to drive robust neural activity in V1.

**Fig. 3.**
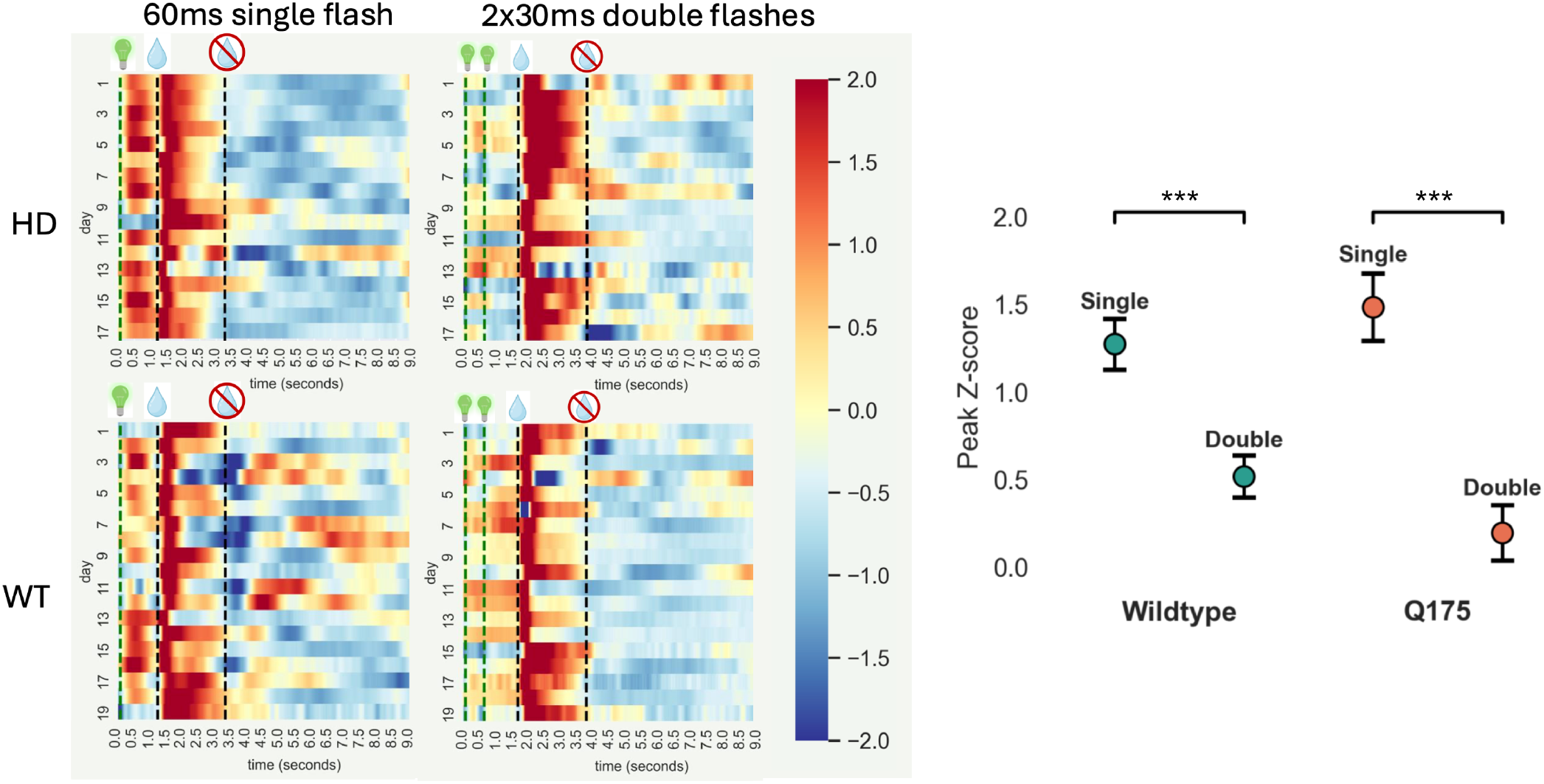
Visual stimulus duration analysis. (Left) Heatmaps of functionally defined primary visual region activity averaged by day and across mice (n=4). The green line at time 0 represents the flash onset. In the double flash condition, the second flash occurs 0.5 seconds later. Spouts move in 1.2 seconds after the final flash. (Right) Visual stimulus duration peak analysis. Peak Z-score value in V1 within 1.2 seconds after flash onset. A significant difference (P *<* 0.0001) was observed in both wild-type and HD mice, with the shorter 30-ms double-flash condition eliciting a weaker cortical response compared to the longer 60-ms single-flash condition.

In contrast, the single 60-ms flash condition elicited a much stronger and more consistent response in the primary visual cortex. On most experimental days, this condition produced a clear peak in cortical activity immediately following stimulus presentation, indicating that the visual system was effectively detecting and processing the longer flash duration. The disparity between the two conditions suggested that stimulus duration played a critical role in evoking a measurable neural response in V1.

To quantitatively assess differences in peak cortical responses between flash conditions, the peak activity for WT mice during the single-flash condition was extracted across all days, and the average response was calculated within 1.2 seconds following stimulus onset.

This was then compared to the peak activity of the double-flash condition (Figure 3). An independent t-test statistical analysis, with a Bonferroni multiple comparisons correction, revealed a significant difference between the two conditions in WT mice (p < 0.0001), confirming that the single-flash stimulus generated a stronger cortical response. The same analysis was conducted for HD mice, where the difference between the single and double-flash conditions was also statistically significant (p < 0.0001). These findings suggest that the reduced response to the double-flash condition was consistent across genotypes.

### Cortical activity differences between successful lick and no-lick trials

As shown in Figure 2 (bottom panel), a mouse cohort (n=5) underwent Stage 2, in which a single spout was presented along with its corresponding stimulus – either a 120 ms single flash of light paired with the left spout or two 60ms flashes paired with the right spout – for a water reward. Cortical imaging results from this cohort were futher analyzed. The double flash condition is included in the Supplementary Figures, but the primary focus of this analysis is on the single-flash condition.

Each trial was categorized based on whether the mice successfully licked the spout. Layer 2/3 cortical calcium responses were then averaged across all trials within each daily session for individual mice, followed by averaging across genotypes (WT and HD). The last three days were averaged to assess the cortical data, as there was an equal distribution of successful and no-lick trials.

When plotting the time course of cortical activity from stimulus onset to the movement of the spouts, both lick and no-lick trials exhibited a visual cortex response. As expected, successful lick trials (Figure 4) evoked a significantly larger motor cortex response following spout movement, likely due to licking activity. This response extended into the barrel fields and other associated sensory regions.

**Fig. 4.**
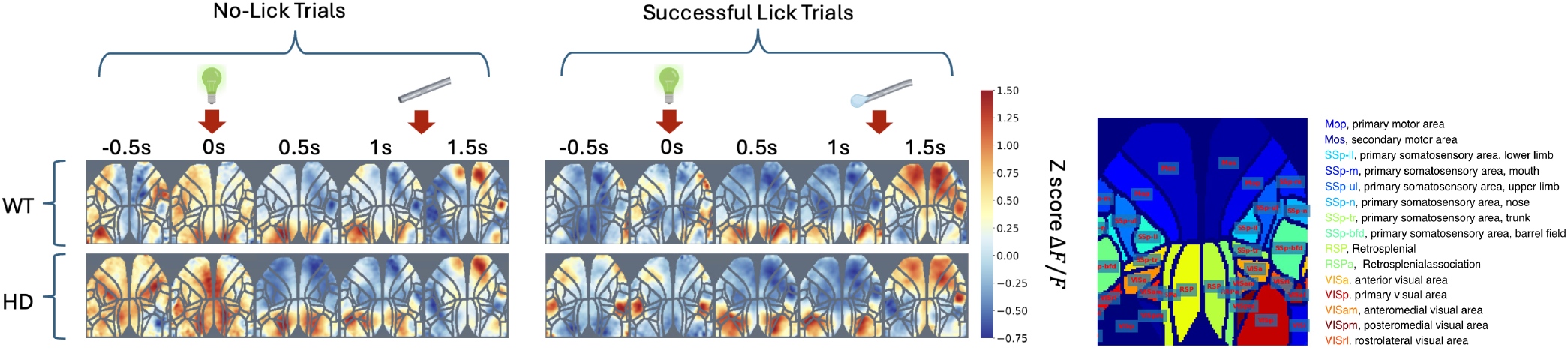
(Left) Single 120ms flash stimuli, paired with left spout presentation. Cortical window activity was average across trial within a session and across genotypes. No-lick condition were trials where the mice did not lick the spout, whereas successful lick conditions were the mice licked. Time zero is the light presentation, the second arrow represents the spout/reward presentation. (Right) Allen Brain Atlas: Mouse Brain regions segmented

This pattern was consistently observed across both WT and HD mice, as well as under both single and double flash conditions (Figure 4, Figure S2). To further investigate this effect, the cortical window was segmented into distinct brain regions based on the Allen Brain Atlas (Figure 4), allowing for a more detailed analysis of regional activity. Following spout movement, a pronounced increase in cortical activity was evident in motor regions, illustrated in Figure 5. This effect is further quantified in Figure 6, where activity traces reveal a sharp spike in motor cortical regions during successful lick trials, whereas no-lick trials exhibit a muted response.

**Fig. 5.**
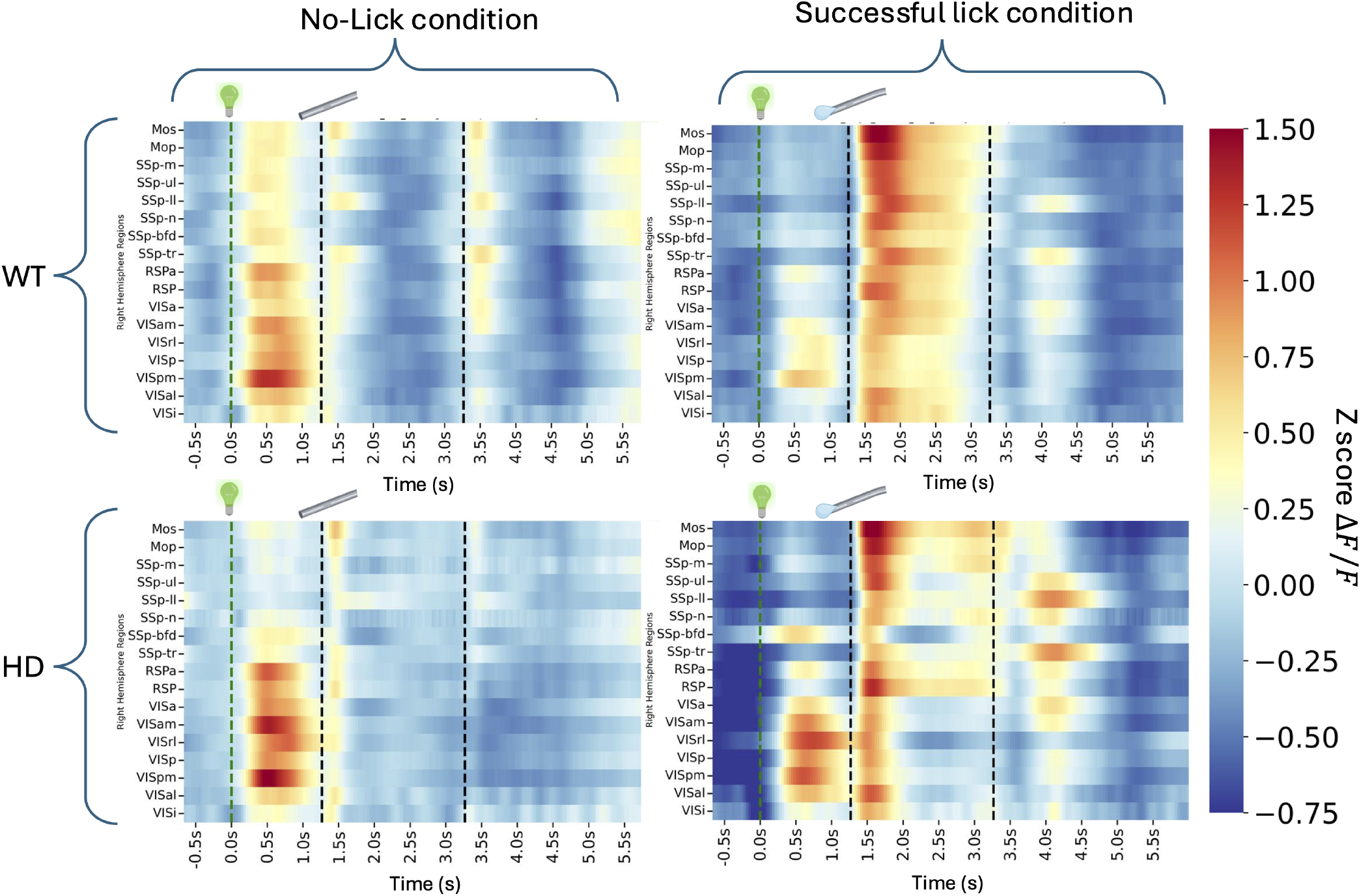
Heatmaps of cortical activity segmented by Allen Brain Atlas regions. 120ms Single flash condition, with left spout presentation. Panels are all in the right hemisphere. Trials were split based on no-lick or successful lick of the mouse in response to the reward presentation post visual stimuli.

**Fig. 6.**
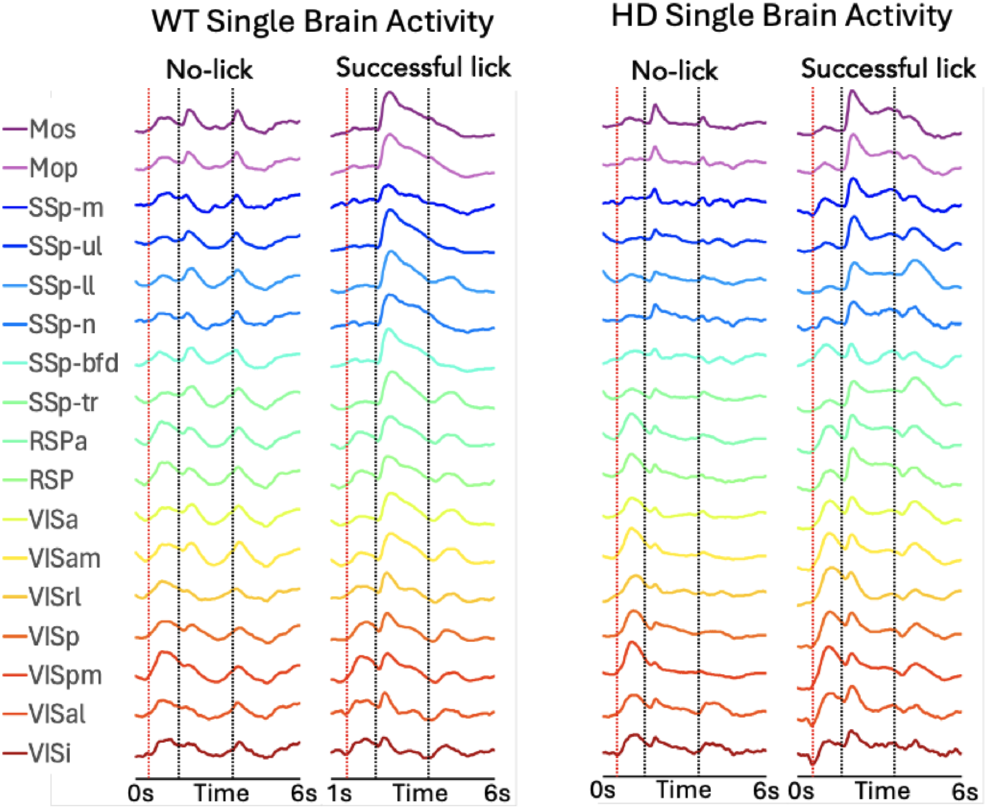
Brain region activity traces across different conditions. The first vertical line (red) is the flash onset, the first black line is the spout presentation, the second black line is the spout removal.

### Increased cortical activity preceding spout movement in no-lick versus successful lick condition

By observing the cortical activity in the different conditions, an interesting distinction emerged between successful and no-lick trials; no-lick trials exhibited heightened cortical activity before the visual stimulus onset at −0.5s (Figure 4). This pattern was also present in the double flash condition (Figure S2), where cortical activity was already elevated at time 0s and continued increasing at 1s. This suggests that when mice were less attentive to the visual stimulus, their cortical activity was more distributed across unrelated brain regions, potentially leading to failures in recognizing and responding to the spout reward. The heatmaps further illustrate this phenomenon—successful lick trials showed notably reduced baseline activity before flash onset, whereas no-lick conditions exhibited more robust activity across all brain regions. While both WT and HD mice showed strong activation in the visual cortex and associated regions regardless of trial outcome, the key difference emerged in non-visual regions. In no-lick trials, motor and sensory areas displayed activity slightly above baseline (.6.F/F z-score > 0), whereas in successful lick trials, these regions were notably suppressed, with z-scores falling mostly below 0. This suggests that successful trials involve more precise encoding of visual stimuli, potentially enabling a co-ordinated motor response, whereas failed trials may reflect inefficient sensorimotor integration or impaired cortical motor planning.

### Visual responses in HD have a higher peak compared to WT

Following the visual stimulus, the visual cortex and its associated regions are activated, as expected, to process the sensory input. This response is consistently observed across both the 120 ms single-flash and the two 60 ms double-flash conditions, as well as in both successful and no-lick trials. Notably, while visual cortex activation is evident in both WT and HD mice, HD mice exhibit an amplified response compared to their WT counterparts.

This heightened response is apparent across multiple forms of analysis. In the heatmaps (Figure 5), HD mice display a more pronounced visual activation compared to WT. Similarly, the traces in Figure 6 show that HD mice exhibit a sharp, rapid peak in visual activity following the stimulus across both no-lick and successful lick trials, with consistently higher amplitudes than WT mice, whose responses are notably attenuated across all visual regions. The convergence of these findings across different visualization methods suggests that HD mice exhibit exaggerated cortical responses to visual stimuli.

### Sustained motor cortex activity following spout presentation in HD mice compared to gradual decrease in WT

When examining motor cortex activity in secondary motor (Mos or M2) and primary motor cortex (Mop or M1) (Figure 6), WT mice exhibit a higher peak response immediately following spout presentation compared to HD mice. After reaching this peak, WT activity gradually declines to baseline over *>*3 seconds in a time-dependent monotonic manner. In contrast, HD mice show a different morphology of decay, with a 2 phase, rapid decline followed by a plateau phase that then declines to baseline after *>*3.5 seconds. This sustained activation is not limited to the motor cortex but is also observed in other cortical regions, where HD mice maintain a heightened level of activity until the spouts move out.

## Discussion

This study provides an exploratory analysis of cortical activity in WT and HD mice during a cue-based discrimination task, leveraging simultaneous mesoscale imaging and behavioral readouts. As part of this analysis, we found that visual stimuli needed to be at least 30–60 ms in duration to robustly activate the visual cortex in the context of a paired sensory-reward cue. This suggests a temporal threshold for reliable cortical engagement, indicating that shorter stimuli may not sufficiently drive perceptual processing or support the formation of sensory-reward associations (19, 20).

Consistent with prior reports, HD mice exhibited heightened visual cortex responses compared to WT, suggesting differences in sensory processing that may reflect hyperexcitability or altered sensory integration (11, 16, 21). This may also reflect broader disruptions in cortical regulation, potentially related to aberrant circuit activation previously observed in HD models. Although this pilot study only encompassed the training stages leading up to a two-alternative forced choice task, the custom-designed rig was built specifically to probe the relationship between these exaggerated sensory responses and behavioral outcomes. Notably, even at this early stage, differences in cortical dynamics between HD and WT mice were evident, underscoring this platform’s potential for future in-depth investigations.

Task performance also influenced cortical activity patterns. Successful lick trials were associated with stronger motor cortex activation following spout presentation, whereas no-lick trials displayed increased pre-stimulus activity and reduced motor engagement. These patterns are consistent with expectations, as active licking involves coordinated motor output, validating that the imaging data reflect task-relevant neural processes (22). Interestingly, WT mice showed a sharp cortical motor activation peak followed by a monotonic decline, while HD mice displayed a more sustained cortical motor response. This persistent activation could reflect impaired inhibition or excessive recurrent activity, both features implicated in HD-related motor dysfunction (12). Additionally, the lower overall cortical motor peak in HD mice may suggest reduced motor drive or less efficient motor execution (23).

Beyond motor output, differences in cortical activation patterns during reward-predictive trials offer insight into how learning may shape sensory processing. Previous studies have shown that expert performance is characterized not only by stronger activation in relevant regions but also by a suppression of activity in non-task-related areas (24). Similarly, our data revealed that successful lick trials featured more selective activation in visual areas, whereas no-lick trials were marked by widespread cortical activation preceding reward presentation. This suggests that effective learning may rely on both enhancement of relevant signals and suppression of competing inputs, promoting more precise visual processing (25). With a sufficiently sensitive decoder, these pre-response dynamics could potentially predict behavioral outcomes, making this an intriguing direction for future analysis (26).

While these findings remain observational and should be interpreted cautiously due to the small sample size, they provide an important first look at how cortical circuits differ between HD and WT mice during active behavior. Continued work using this refined behavioral paradigm with larger cohorts will be essential to confirm these initial observations and further dissect the cortical mechanisms underlying sensory-guided decision-making in HD.

## AUTHOR CONTRIBUTIONS

KT, DR, JM, and LAR designed the experiments. The mesoscale camera was designed and developed by THM. KT performed the research. KT analyzed the data, and DR, JM, THM, and LAR contributed to data interpretation. KT drafted the manuscript with input and editing from THM and LAR.

## ACKNOWLEDGEMENTS

This work was supported by resources made available through the NeuroImaging and NeuroComputation Centre at the Djavad Mowafaghian Centre for Brain Health (RRID: SCR_019086); Canadian Institutes of Health Research (CIHR) Grant PJT-191742 to LAR. Foundation Grant FDN-143209, PJT-180631, and the National Science and Engineering Council of Canada (NSERC) Grant GPIN-2022-03723 to THM. CIHR CGS-M award to KT.

We thank Lily Zhang for technical genotyping and cranial window assistance.

## Supplementary Material 1: Mouse cranial window surgery SOP

### Headbar and Coverslip Preparation

1. Cut a coverslip into a 1 cm x 1 cm square (or to fit the size of the headbar window) using a diamond cutter.
2. Autoclave headbar and coverslip *Mouse Preparation*
3. Measure the weight of the mouse.
4. Induce anesthesia using an isoflurane induction chamber (5 on the anesthesia machine scale for induction, then maintain at 2).
5. Insert a temperature sensor into the rectum to maintain the feedback-controlled heating pad (WPI) at 37°C.
6. Apply Refresh™ Lacri-Lube ophthalmic ointment to the eyes for hydration.
7. Shave the top of the head, extending down to the neck.
8. Place the mouse under the microscope and maintain anesthesia. *Surgical Procedure*
9. Clean the shaved area by applying iodine followed by 70% acetone.
10. Administer the following subcutaneous (SQ) injections: - 0.3 ml of 0.9% saline to prevent dehydration during surgery. - 5 mg/kg meloxicam. - 7 mg/kg bupivacaine at surgery local area
11. Carefully remove all skin from the shaved area (from the top of the head to the neck).
12. Apply 3M Vetbond tissue adhesive along the edges of the cut area to prevent tissue fluid exudation.
13. Use Krazy Glue to attach the edge of the headbar in place. Hold it for 5 minutes to allow the glue to settle. *Dental Cement and Coverslip Placement*
14. In a tray, prepare the dental cement as follows: - Mix Parkell, C&B Metabond Quick Base (solvent) with Parkell, C&B Metabond Clear Powder (powder). - Add Parkell, C&B Amalgambond Universal Catalyst to the mixture to form the cement.
15. Immediately apply the dental cement to the cutout area and place the prepared glass coverslip on top. Press gently to eliminate air bubbles under the coverslip.
16. Wait 30 minutes for the dental cement to set.
17. Add additional dental cement around the headbar and the base of the skull to secure the structure and to ensure no any tissue exposure in the air. Allow 30 minutes to dry. *Post-Surgical Care*
18. Place the mouse in a recovery cage with a heating pad set to 37°C. Once the mouse is fully awake and moving, return it to the home cage.
19. Administer a second SQ injection of 5 mg/kg meloxicam 10-12hr after surgery.
20. Monitor the mouse’s weight for 3 days or until weight stabilizes.

## Supplementary Material 2: Figures

**Fig. S1.**
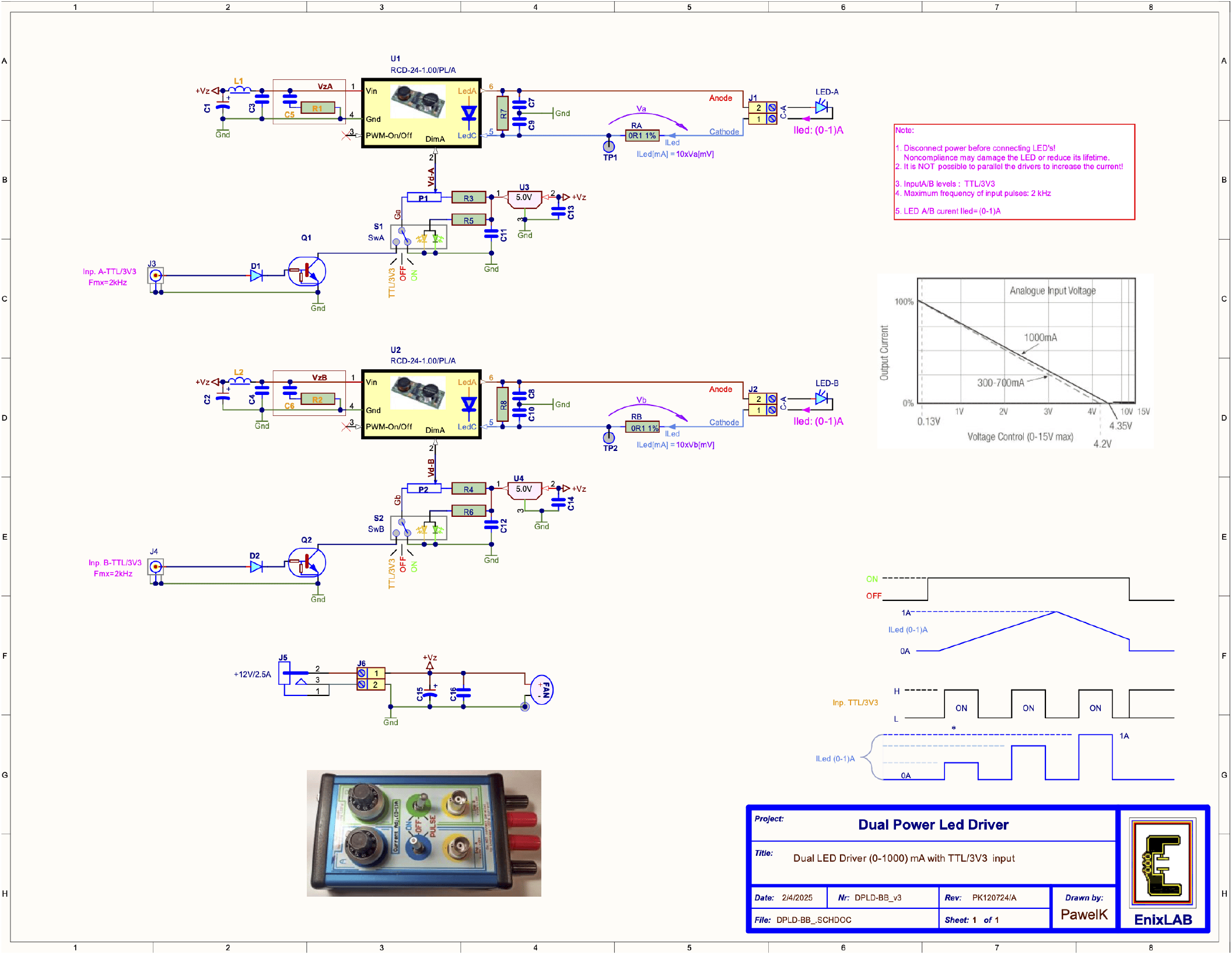
Custom dual power LED driver schematic from the EnixLAB in Vancouver, BC, Canada

**Fig. S2.**
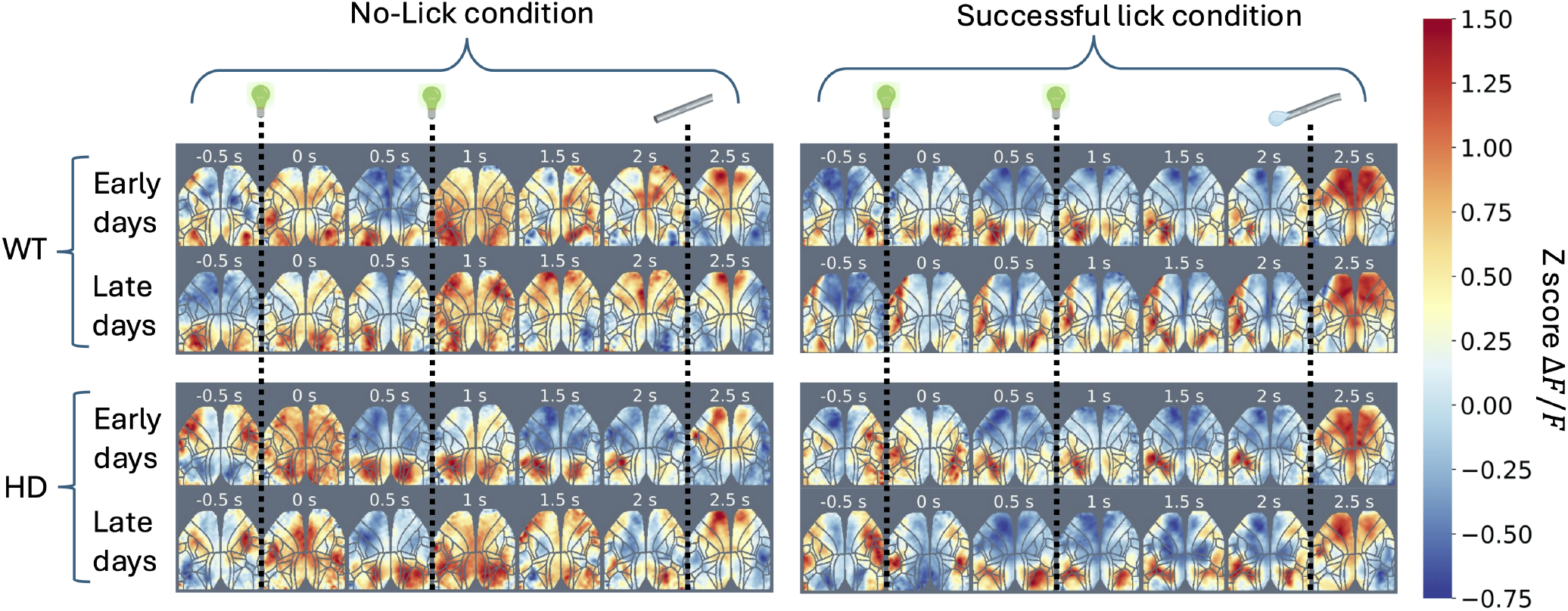
Double flash condition. Mice were presented with two 60ms pulses of light separated by 1 second. The right spout was moved as the water reward. This condition showed similar visual activity post visual stimulus and motor cortex activation on successful lick conditions compared to no-lick conditions. First flash of light occurs at 0 seconds, followed by the second flash shortly after 1 second. The spouts presentation are a little past 2 seconds.

**Fig. S3.**
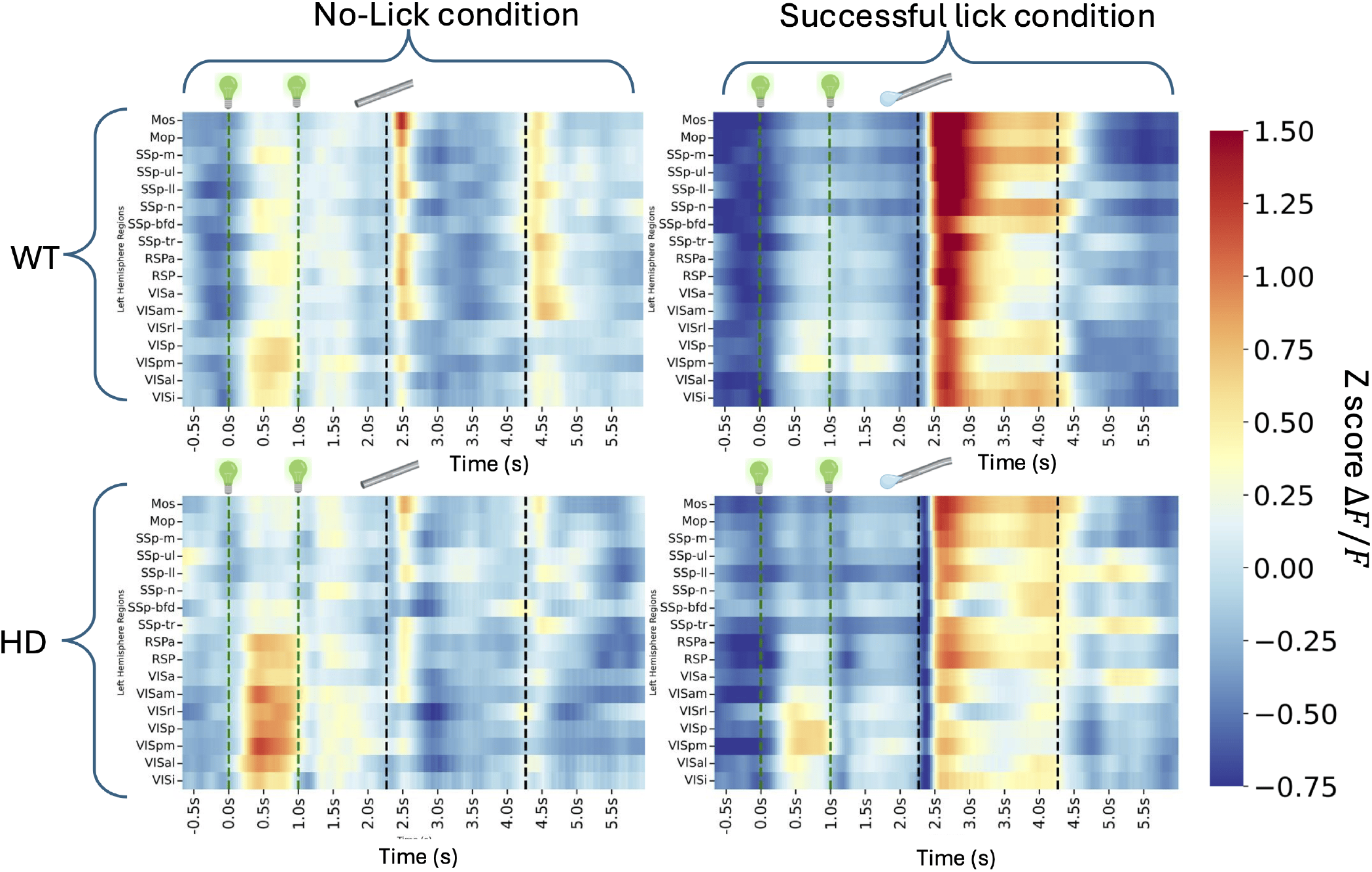
Left hemisphere, double flash heatmaps of brain region activity

